# Temporal Dysbiosis of Infant Nasal Microbiota Relative to Respiratory Syncytial Virus Infection

**DOI:** 10.1101/2020.04.30.071258

**Authors:** Alex Grier, Ann L. Gill, Haeja A. Kessler, Anthony Corbett, Sanjukta Bandyopadhyay, James Java, Jeanne Holden-Wiltse, Ann R. Falsey, David J. Topham, Thomas J. Mariani, Mary T. Caserta, Edward E. Walsh, Steven R. Gill

**Affiliations:** Department of Microbiology and Immunology, University of Rochester School of Medicine and Dentistry, Rochester, NY, USA; Department of Biostatistics and Computational Biology, University of Rochester School of Medicine and Dentistry, Rochester, NY, USA; Division of Neonatology and Pediatric Molecular and Personalized Medicine Program, University of Rochester School of Medicine and Dentistry, Rochester, NY, USA; Division of Pediatric Infectious Diseases, University of Rochester School of Medicine and Dentistry, Rochester, NY, USA; Genomics Research Center, University of Rochester School of Medicine and Dentistry, Rochester, NY, USA; Department of Medicine, Rochester General Hospital, University of Rochester School of Medicine and Dentistry, Rochester, NY, USA

**Author notes:** **Address for Correspondence:** Steven R. Gill, PhD, Department of Microbiology and Immunology, University of Rochester School of Medicine and Dentistry, 601 Elmwood Avenue, Rochester, NY, 16462, USA, Rochester NY, 14642, USA. **DECLARATIONS** **Ethics approval and consent to participate** Written informed consent was obtained from parent or guardian of all participating infants. The institutional review board at the University of Rochester School of Medicine and Strong Memorial Hospital approved the study. **Presentation of information from this manuscript at meetings** Data from this manuscript has not been presented at meetings outside of University of Rochester. **Permission for personal communications** We grant permission for all personal communications with the Journal of Infectious Diseases. **Competing interests** The authors declare that they have no competing interests. **Funding** This project has been funded in whole or in part with Federal funds from the National Institute of Allergy and Infectious Diseases, National Institutes of Health, Department of Health and Human Services, under Contract No. HHSN272201200005C.

**Keywords:** microbiota, RSV, infant respiratory disease

## Abstract

**Rationale:** Respiratory Syncytial Virus (RSV) infection is a leading cause of infant respiratory disease and hospitalization. Infant airway microbiota occupying the nasopharynx have been associated with respiratory disease risk and severity. The extent to which interactions between RSV and microbiota occur in the airway, and their impact on respiratory disease severity and infection susceptibility, are not well understood.

**Objectives:** To characterize associations between the nasal microbiota and RSV infection before, during, and after infants’ first respiratory illness.

**Methods:** Nasal 16S rRNA microbial community profiling of two cohorts of infants in the first year of life: 1) a cross-sectional cohort of 89 RSV infected infants sampled during illness and 102 population matched healthy controls, and 2) an individually matched longitudinal cohort of 12 infants who developed RSV infection and 12 who did not, sampled at time points before, during, and after infection.

**Measurements and Main Results:** We identified 12 taxa significantly associated with RSV infection. All 12 were differentially abundant during infection, with seven differentially abundant prior to infection, and eight differentially abundant after infection. Eight of these taxa were associated with disease severity. Nasal microbiota composition was more discriminative of healthy vs. infected than of disease severity.

**Conclusions:** Our findings elucidate the chronology of nasal microbiota dysbiosis and suggest an altered developmental trajectory associated with first-time RSV infection. Microbial temporal dynamics reveal indicators of disease risk, correlates of illness and severity, and the impact of RSV infection on microbiota composition. Identified taxa represent appealing targets for additional translationally-oriented research.

## Introduction

The composition and function of host-associated microbial communities are associated with many aspects of health and disease [1]. These relationships between the microbiome and host biology exhibit spatial and temporal dependencies, with relevant interactions manifest by the microbiota of specific body sites during critical periods of host development, environmental exposure, pathogenesis, illness, or convalescence [2–6]. Specifically, there is a growing body of evidence that the microbiome influences immune maturation and function [7–9], mucosal surface physiology [10, 11], and the risk and severity of acute and chronic respiratory diseases [12–16].

Respiratory Syncytial Virus (RSV) is the most significant respiratory tract infection affecting infants. It is the most frequent cause of acute lower respiratory infections in children under five, and a common cause of hospitalization in children under two [17–19]. Approximately one-half of infants are infected with RSV during their first year of life, and nearly all have been infected by two years of age. Severe disease requiring hospitalization occurs in 1-3% of those infected, and in most cases is not accompanied by any of the known risk factors such as age at infection, pre-term birth, underlying cardiopulmonary disease or immunosuppression [20–22]. Additionally, RSV infection in early life has been linked to subsequent development of asthma and chronic obstructive lung disease [19, 23].

Recent studies have identified associations between nasopharyngeal microbiota and RSV clinical manifestations including severity [24, 25]. Nasopharyngeal microbiota composition has been shown to be altered during periods of acute RSV infection and the abundance of certain bacterial taxa have been associated with immune response and disease severity [13, 24–26]. While these findings suggest that respiratory microbiota may play an important role in RSV infection, the spatial and temporal scope of a relationship remains unclear. Specifically, whether associations between RSV infection and microbiota composition are limited to the nasopharynx, and in what sequence and duration they manifest, are not well understood [27, 28].

Here, we analyze the nasal microbiota of two cohorts of infants to elucidate the relationship between airway microbial communities and RSV infection. We used a large cross-sectional cohort of infants comprised of an RSV infected case group sampled during acute illness and a matching healthy control group to characterize the nasal microbiota of acute RSV infection and identify associations with disease severity. To assess associations with the nasal microbiota that may exist before or after RSV infection, we used a smaller longitudinal cohort comprised of a group of infants that developed RSV infection during their first year of life and another group that did not, with each group sampled at matching time points corresponding to before, during, and after acute illness.

## METHODS

### Clinical methods

All study procedures were approved by the University of Rochester Medical Center (URMC) Research Subjects Internal Review Board (IRB) (Protocol # RPRC00045470) and all subjects’ caregivers provided informed consent. The infants included in the study were from the University of Rochester Respiratory Pathogens Research Center AsPIRES [29] and PRISM studies and cared for prior to discharge in the URMC Golisano Children’s Hospital and Rochester General Hospital newborn nurseries and birthing centers. For the cross-sectional cohort (**Table 1A**), we analyzed 191 nasal samples from 89 subjects with acute RSV infection and 102 healthy subjects. Control samples and subjects were selected to minimize population level differences in age at the time of sampling, gestational age at birth, and mode of delivery. For the longitudinal cohort (**Table 1B**), we collected 72 nasal samples from 12 RSV positive subjects and 12 healthy subjects. Samples were collected from the RSV group at approximately one month of age, during acute RSV infection, and approximately one month after illness, and at corresponding timepoints from the healthy controls. Control subjects were selected to match on an individual basis by sex, mode of delivery, and gestational age at birth, and samples were selected to match by age. Subjects were eligible as controls if they had no respiratory illness between birth and at least ten days after the last sample. Patient metadata for the cross-sectional and longitudinal cohorts is included in the online **Supplemental Table 1.**

**Table 1A.** Summary Characteristics of Cross-Sectional Cohort.

| **Variables**<br>**(mean ± SD or N)** | **Aggregate**<br>**(N=191)** | **Control**<br>**(N=102)** | **Case**<br>**(N=89)** |
| --- | --- | --- | --- |
| Sex (Male/Female) | 105/86 | 61/41 | 44/45 |
| Mode of Delivery (Vaginal/C-section) | 128/63 | 68/34 | 60/29 |
| Gestational Age at Birth (Weeks) | 39.23 ± 1.17 | 39.27 ± 1.13 | 39.18 ± 1.23 |
| Age at Sampling (Months) | 3.09 ± 2.21 | 3.01 ± 2.19 | 3.18 ± 2.24 |
| Any Antibiotics To Date | 0 | 0 | 21 |
| Severity Score | 2.24 ± 3.01 | 0 ± 0 | 4.81 ± 2.65 |
| Severity Group (Mild/Severe) | NA | NA | 30/59 |

**Table 1B.**
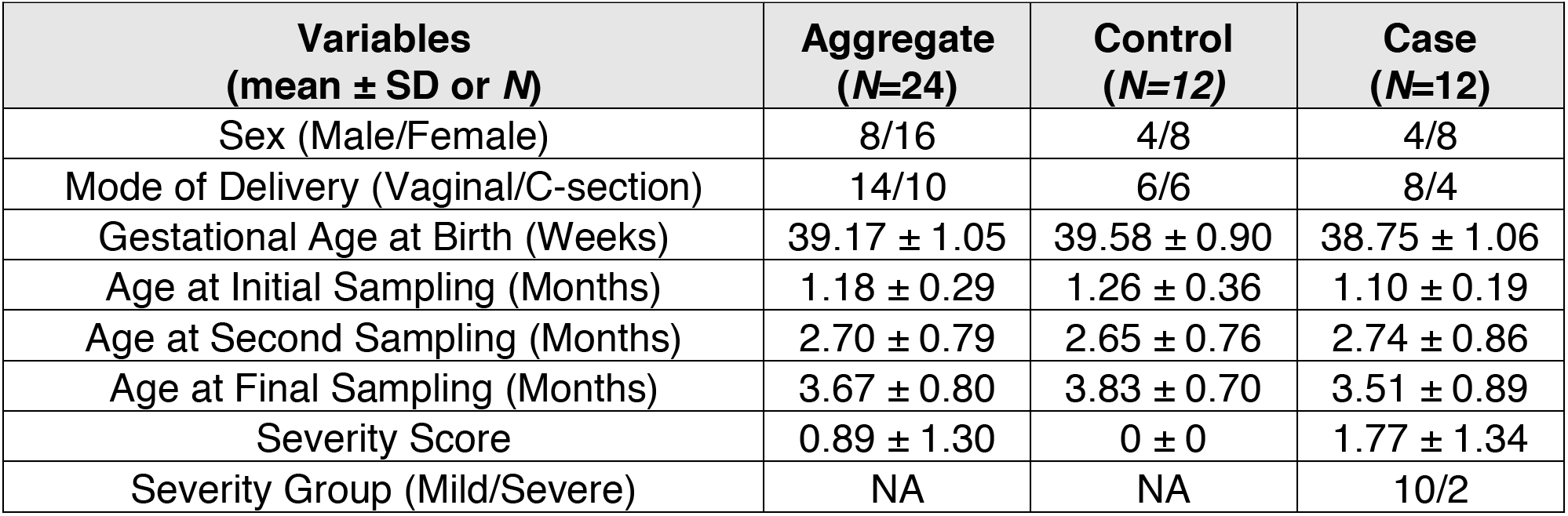
Summary Characteristics of Longitudinal Cohort.

### 16s rRNA amplicon sequencing

Genomic DNA was extracted and the V1-V3 16S rRNA hypervariable region was sequenced as described previously [4]. Bioinformatics processing was performed with QIIME 2 [30], using DADA2 [31] for denoising and the GreenGenes reference database [32, 33] as the basis of taxonomic classification. Additional methodological details of sample preparation, sequencing, controls, and bioinformatic processing are available in **Supplemental Methods**.

### Associations of taxon abundance with RSV infection and disease severity

Because differential abundance testing of high-throughput sequencing-based microbial community profiling data is relatively immature, with no consensus methodology [34, 35], we applied four prominent univariate and multivariate algorithms which were selected to be complementary in terms of their strengths and technical limitations. We required that significant results be corroborated across multiple methods to be accepted. Details of diversity analyses and machine learning classification/regression analyses are available in **Supplemental Methods**.

## RESULTS

### Overview of infant cohort

The cross-sectional case-control cohort yielded 191 nasal samples with 16S rRNA sequencing data from 89 subjects with acute RSV infection and 102 matched healthy subjects **(Table 1A)**. The average number of reads per sample was 64,320 with 180 samples having at least 5,000 reads. All subjects were full-term and less than 10 months of age, and the ill and healthy groups matched at the population level in terms of sex, gestational age at birth, mode of delivery, and age at the time of sampling. Infected subjects were divided into mild and severe based on a threshold Global Respiratory Severity Score (GRSS) of 3.5, yielding groups of 30 and 59, respectively [36]. Severity scores and additional patient metadata for the cross-sectional and longitudinal cohorts are in **Supplemental Table 1**.

The longitudinal cohort yielded 72 nasal samples with 16S rRNA sequencing data corresponding to 12 healthy controls and 12 RSV positive subjects sampled at three time points: one month of age, during acute illness (and corresponding age for healthy controls), and one month after illness **(Table 1B)**. The average number of reads per sample was 47,745, with 67 samples having at least 5,000 reads. Healthy controls closely matched RSV positive subjects in terms of sex, mode of delivery, gestational age at birth, and age at the time of sampling. All subjects were full term and less than one year of age. Healthy controls did not develop symptomatic respiratory infection between birth and at least 10 days after their last sample was taken. Notably, only two of the RSV cases in this cohort exhibited severe disease (GRSS > 3.5).

### Microbiota diversity and associations with RSV infection and severity

In the cross-sectional cohort, alpha diversity as measured by Faith’s index was elevated in RSV positive subjects at the time of infection relative to age matched healthy controls (p = 0.039), with a greater difference observed in the group of subjects with severe disease (mean Faith’s index of healthy = 1.933, mild illness = 2.176, severe illness = 2.250). The difference between subjects with mild and severe infection was not significant, however, the correlation coefficient between severity score and Faith’s index was positive (*r* = 0.134; p=0.065). There were no significant differences in alpha diversity as measured by the Shannon index, suggesting that the observed differences reflect increased phylogenetic heterogeneity in the subjects with infection, as opposed to a greater number of total species or more even distributions of species’ relative abundances.

In the longitudinal cohort, Weighted and Unweighted Unifrac distances were used to assess beta diversity at each visit, and to assess the magnitude of change within individuals from visit to visit **(Figure 1A)**. At all three time points, significant differences were found between the group that developed RSV infection and the group that did not, based on the Weighted Unifrac metric (initial visit p = 0.032, illness and age matched healthy visit p = 0.009, follow-up visit p = 0.012). By Unweighted Unifrac, these differences were significant at the initial visit (p = 0.035) and the illness visit (p = 0.011), and approached significance at the post-illness visit (p=0.078). By both metrics, the largest, most significant difference was observed at the illness visit (and the corresponding timepoint for the healthy controls). Further examination of beta diversity during illness using the cross-sectional cohort **(Figure 1B)** revealed more significant differences between healthy subjects and severely ill subjects (Unweighted Unifrac p = 0.003, Weighted Unifrac p = 0.003) than between healthy subjects and subjects with mild disease (Unweighted Unifrac p = 0.036, Weighted Unifrac p = 0.005), as well as greater differences between healthy and RSV infected infants when the infection occurred at younger ages (among subject 0-3 months old, Unifrac PERMANOVA healthy vs. mildly ill p = 0.538 (Unweighted) and 0.084 (Weighted), healthy vs. severely ill p = 0.001 (Unweighted) and 0.001 (Weighted); among subjects > 6 months old, healthy vs mildly ill p = 0.931 (Unweighted) and 0.191 (Weighted), healthy vs. severely ill p = 0.309 (Unweighted) and 0.389 (Weighted)). Assessing the magnitude of longitudinal changes by the Unweighted Unifrac metric, the within subject change from the initial visit to the illness visit, and the corresponding time point in healthy subjects, was larger among the subjects that developed infection than those that remained healthy (p=0.061). All computed alpha and beta diversity values are in **Supplemental Table 2.**

**Figure 1.**
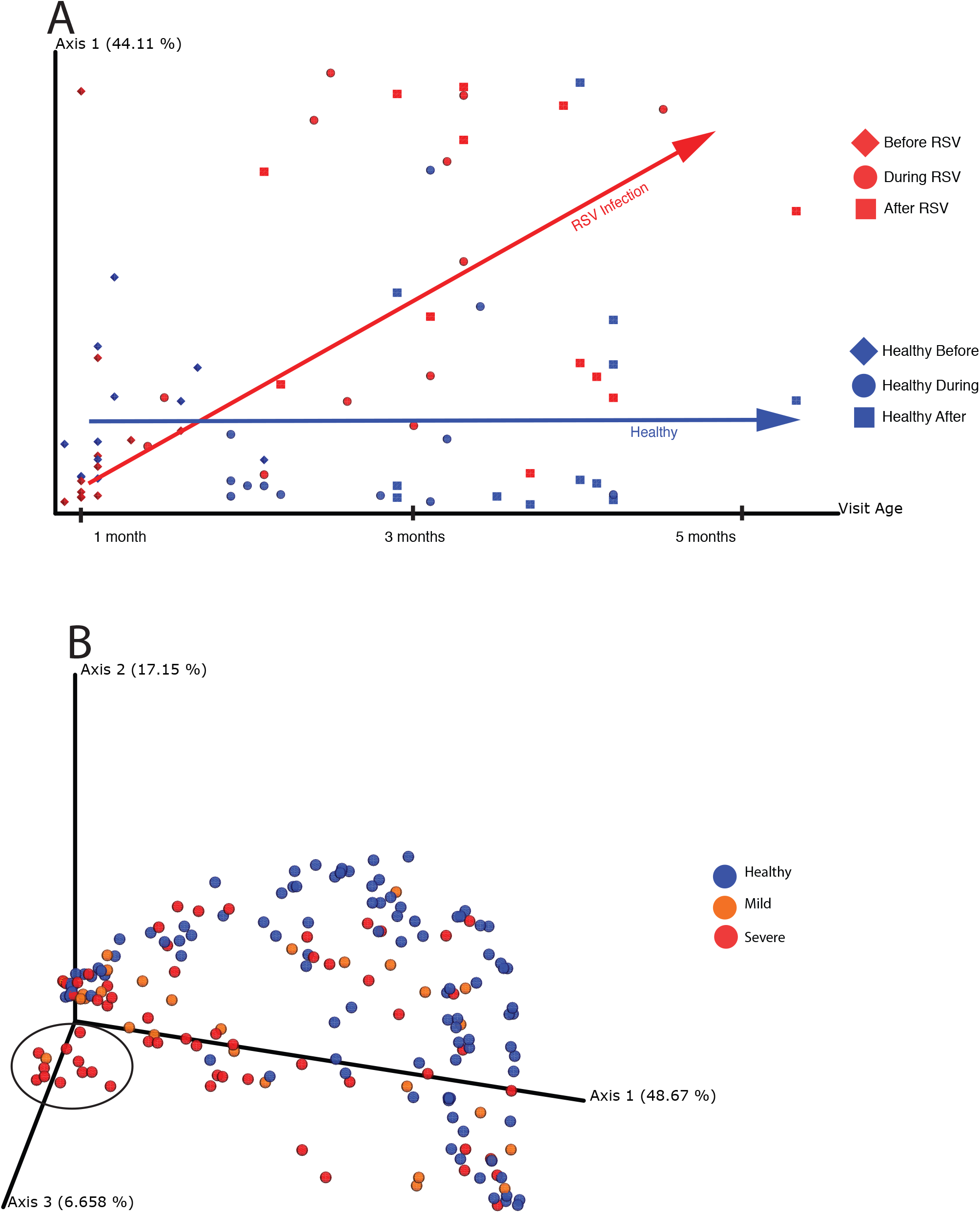
Principal coordinate analysis (PCoA) of weighted Unifrac distances was used to visualize relationships between the nasal microbiota of infants with respect to RSV infection, illness severity, and time. Weighted Unifrac distances quantify the compositional dissimilarity between microbial communities, incorporating information about the phylogenetic relatedness between bacteria observed across samples. PCoA provides a summary representation of overall similarity/dissimilarity relationships among a set of samples, capturing as much information as possible using the fewest number of dimensions/principal coordinates. The proportion of overall variation represented along a single axis is indicated as a percentage in the axis label. **(A)** From the longitudinal cohort only, samples are plotted with principal coordinate one on the y-axis and infant age at the time of sampling on the x-axis. Samples are colored red or blue based on whether or not an infant developed RSV infection (red) at any point during the period of observation, and their shape indicates the time-point at which the sample was taken: initial healthy/pre-illness visit (diamond), illness visit/age-matched healthy visit (circle), or post-illness/age-matched final healthy visit (square). The red and blue arrows indicate observed longitudinal trends within the group of subjects that developed RSV infections and the group that stayed healthy, respectively. **(B)** From the cross-sectional cohort only, samples are plotted in three dimensions using the first three principal coordinates. Samples are colored according to RSV infection status and severity: healthy (blue), mild RSV infection (orange), or severe RSV infection (red). A cluster of subjects in the foreground on the left, notable for dominant abundance of *H. Influenzae*, is circled in black. While no clear segregation is observed between mild and severe illness, healthy samples occupy a notable crescent shaped structure around the illness samples, with the *H. Influenzae* dominated cluster furthest away from this crescent.

### Longitudinal abundance patterns of RSV-associated taxa

The relative abundance of twelve distinct taxa exhibited significant associations with the occurrence of RSV infection according to multiple corroborative statistical assessments. While all twelve taxa were differentially abundant between RSV infected and healthy infants during illness, and the corresponding time point in healthy subjects, they exhibit distinguishable patterns of temporal dynamics, pre- and post-illness occurrence, and associations with illness severity **(Table 2)**. Notably, associations between nasal microbiota and RSV infection are not confined to the period of acute infection: all but one (*Haemophilus*) of the twelve taxa associated with RSV infection are significantly differentially abundant between groups either before or after illness, or both. Most of the taxa (7/12) that are differentially abundant between RSV infected and healthy infants during illness are differentially abundant at the initial visit at one month of age, prior to illness. Similarly, most of the taxa (8/12) that are differentially abundant during illness are differentially abundant after illness. However, persistent differential abundance between groups across all three time points is observed only in a minority (4/12) of taxa. Furthermore, the microbiota differences between health and RSV infection are not simply categorical but vary in magnitude with illness, as most of the taxa (8/12) that are differentially abundant during illness are associated with illness severity. Additionally, most of these severity-associated taxa (6/8) exhibit persistent differences beyond the period of acute illness and are differentially abundant during and after illness, while half (4/8) are differentially abundant prior to illness. Finally, the abundances of most RSV-associated taxa are positively associated with the disease, and only a minority of taxa (5/12) that differ in abundance between groups are elevated in healthy infants.

**Table 2.** Taxa Associated with RSV Infection and Illness Severity and the Time Points at Which They Differ Significantly Between Healthy Infants and Those Who Become Ill with RSV. Member Clades in Parentheses are the Primary Drivers of the Association Indicated in Parentheses.

| **Taxon** | **Positive Association** | **Before Illness** | **During Illness** | **After Illness** | **Illness Severity** |
| --- | --- | --- | --- | --- | --- |
| Alphaproteobacteria | Disease | + | + | + | + |
| Gammaproteobacteria | Disease | + | + | + | + |
| Pseudomonadales (Pseudomonas) | Disease | + | + | + | (+) |
| Moraxella | Disease | + | + | + | - |
| Corynebacterium | Health | + | + | - | - |
| Gluconacetobacter | Disease | + | + | - | + |
| Anaerococcus | Health | + | + | - | - |
| Staphylococcus | Health | - | + | + | + |
| Betaproteobacteria (Burkholderiales) | Disease | - | + | + | (+) |
| Bacilli | Health | - | + | + | + |
| Clostridia | Health | - | + | + | - |
| Haemophilus (influenzae) | Disease | - | + | - | (+) |

*Staphylococcus*, Clostridia (not shown), and Bacilli each exhibit a similar temporal pattern; ubiquitous and comparable in abundance between groups prior to illness, significantly diminished during illness (p <= 0.001, 0.013, & 0.005, respectively) and remain so one month later **(Figure 2A**; p = 0.011, 0.033, & 0.039). The classes Alphaproteobacteria (not shown) and Gammaproteobacteria, and Gammaproteobacteria member clades Pseudomondales (not shown) and *Moraxella*, also exhibit a common pattern in that all four are significantly elevated in the infants that develop RSV infection before (p <= 0.001, 0.033, 0.003, & 0.001), during (p <= 0.036, 0.002, 0.001, & 0.001), and after illness **(Figure 2B**; p = 0.044, 0.009, 0.038, & 0.003). By contrast, *Corynebacterium* and *Anaerococcus* are elevated in infants that do not develop RSV infections at the pre-illness timepoint (p = 0.008 & 0.020, respectively) and the illness timepoint (p < 0.001 & p = 0.008) but do not differ between groups at the post-illness timepoint **(Figure 2C)**. Finally, three taxa exhibit unique temporal trends with respect to illness: Betaproteobacteria increases in abundance over time in the RSV group only **(Figure 2D)** – being significantly elevated during (p = 0.006) and after (p = 0.039) illness – while *Haemophilus* (not shown) is significantly more abundant in the infected group during (p < 0.001) illness and minimally abundant in both groups before and after. *Gluconacetobacter* exhibits a distinct temporal pattern in that it is elevated in the RSV group before (p = 0.003) and during (p < 0.001) illness, but no difference is observed between groups after illness **(Figure 2D)**. The composition of all cross-sectional and longitudinal samples summarized at all taxonomic levels is in **Supplemental Tables 3 and 4.**

**Figure 2.**
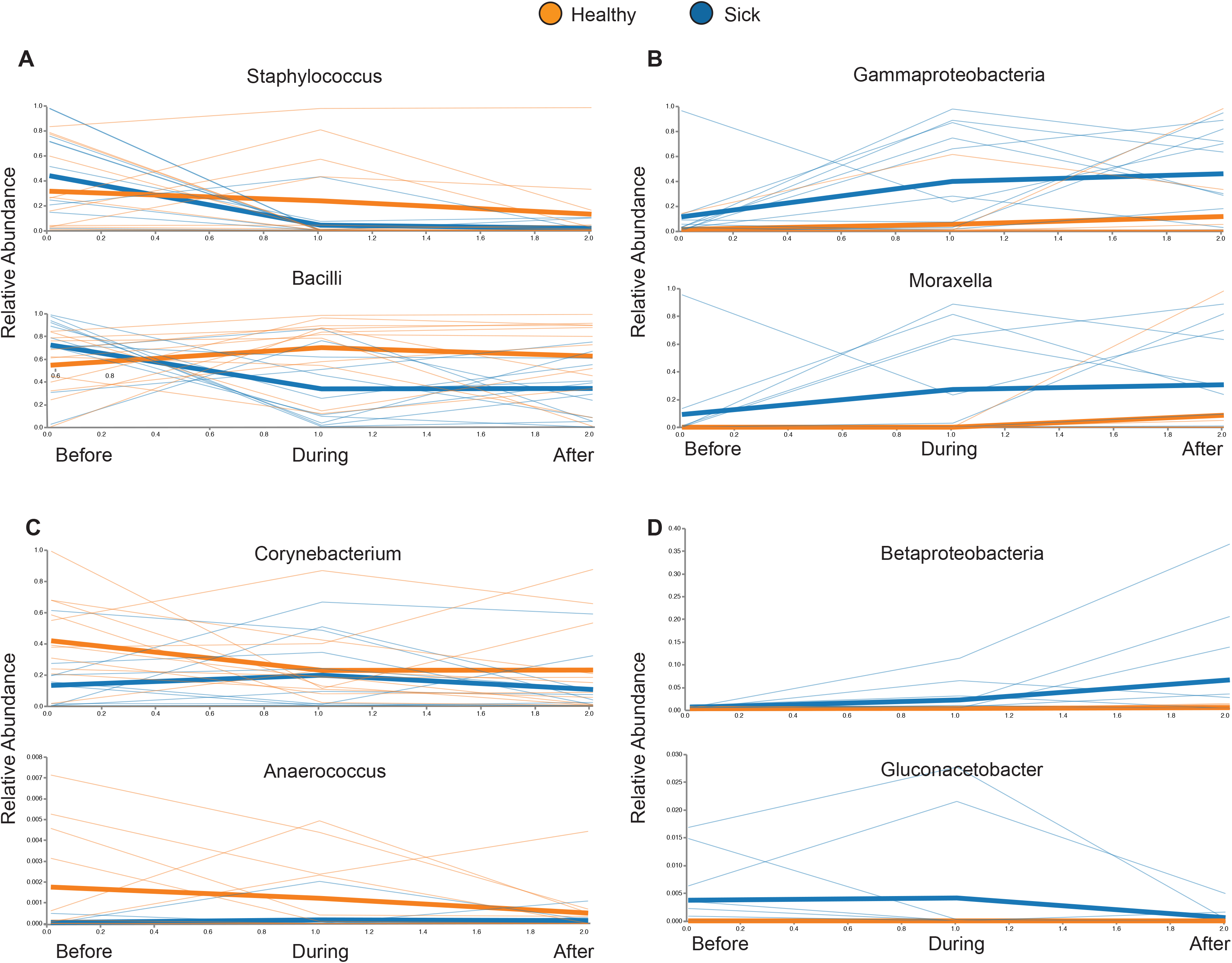
Relative abundances (y-axes) of select taxa at all three time points (x-axes) in the longitudinal cohort. Each thin line corresponds to the abundance of a given taxon in a particular individual, while the thick lines show the mean abundance of each group at each time point. Members of the healthy group are orange and members of the group that developed infection are blue. Significant taxa were grouped based on different temporal patterns of abundance with respect to illness, and each panel shown contains examples from a different group: **(A)** similar abundance between infection and healthy groups prior to illness, but decreased during and after illness in subjects that become infected; **(B)** consistently elevated in the illness group; **(C)** elevated in the healthy group before and during illness, but not after; and **(D)** idiosyncratic temporal dynamics observed in each taxon. Of the members of the fourth group shown here, Betaproteobacteria is nearly absent from all subjects at the pre-illness time point, and then becomes increasingly abundant during and after illness in the infection group while remaining nearly absent from the healthy group. *Gluconacetobacter* is elevated in the infection group prior to and during illness, and substantially diminishes in abundance with convalescence.

### Abundance of taxa associated with severity in acute illness

The abundance of six of the taxa associated with RSV infection are positively associated with severity at the time of acute illness: Alphaproteobacteria (p = 0.026), Gammaproteobacteria (p < 0.001), *Pseudomonas* (p < 0.001), *Gluconacetobacter* (p < 0.001), Burkholderiales (p = 0.015), and *Haemophilus* (p < 0.001), with exceptionally high levels of *Haemophilus influenzae* being very strongly associated (p < 0.001) with severe disease **(Figure 3)**. *Pseudomonas* and Burkholderiales are the primary drivers of associations between severity and their corresponding clades, Pseudomondales and Betaproteobacteria. The abundance of Bacilli (p < 0.001) and *Staphylococcus* (p < 0.001), conversely, are negatively associated with disease severity at the time of illness. As described above, *Moraxella*, *Corynebacterium*, *Anaerococcus*, and Clostridia are associated with the occurrence of RSV infection (or lack thereof), but they are not associated with severity of disease.

**Figure 3.**
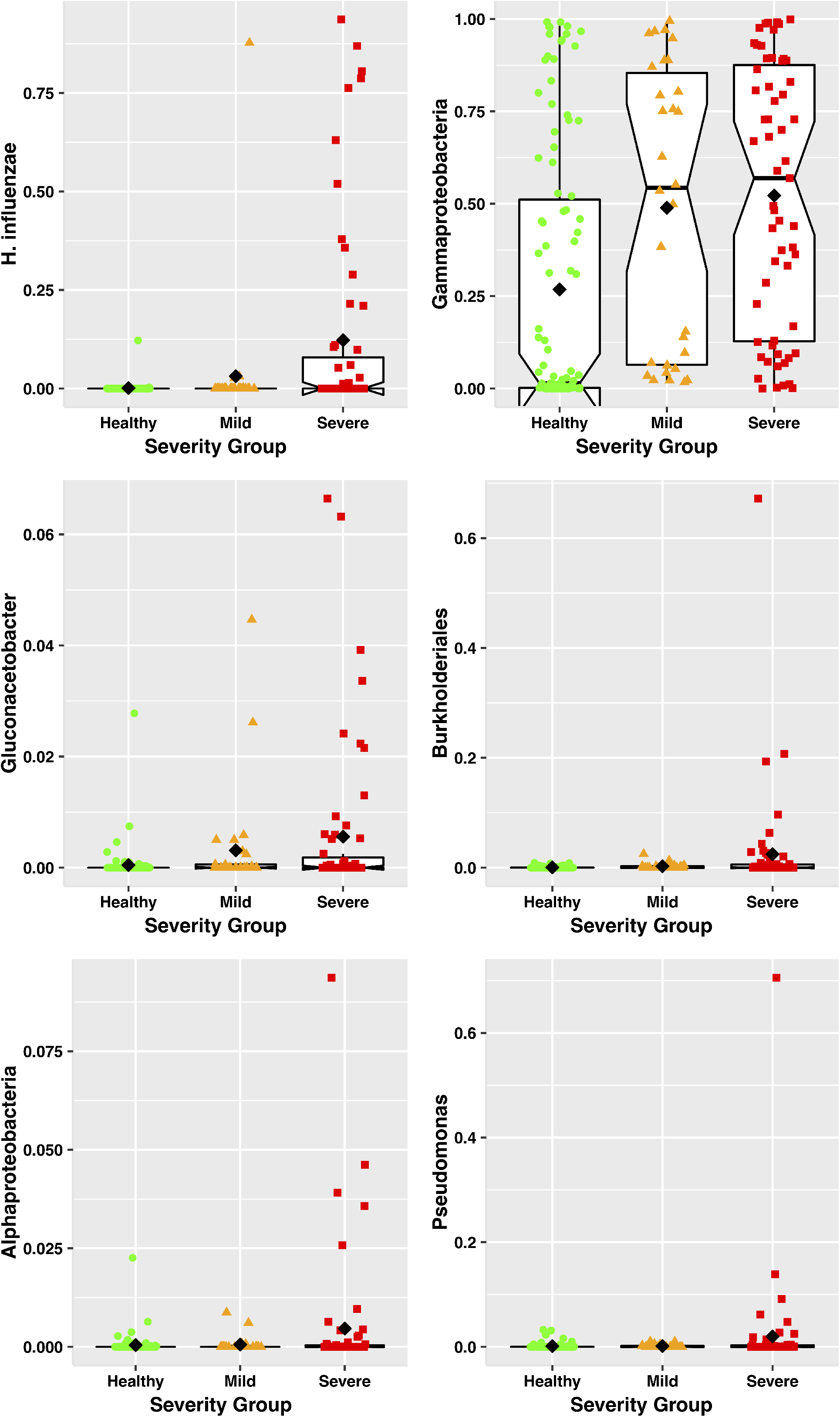
Relative abundances (y-axes) of select taxa significantly associated with more severe disease in the cross-sectional cohort, with samples grouped by dichotomizing illness based on severity into mild and severe groups (x-axes), using a severity score threshold of 3.5. Each colored point represents the relative abundance of a given taxon in a single individual, with columns (left to right), shapes (circle, triangle, square), and colors (green, orange, red) distinguishing between healthy, mild illness, and severe illness groups, respectively. The black diamonds indicate the group mean for each group. Box plots are overlaid on each group, centered on the group median, with notches indicating an approximately 95% confidence interval, boxes indicating boundaries of the first and third quartiles, and whiskers extending to the largest and smallest values no further than 1.5*(inter-quartile range) from the boxes. Points beyond the whiskers are commonly considered outliers, which in this case would suggest that many of the observed associations between taxon relative abundance and illness severity are driven primarily by outliers, or that taxon abundance in severely ill infants comprises more than one underlying distribution.

### Predicting RSV infection status and illness severity from microbiota composition

To further assess the relationship between nasal microbiota and RSV infection using the cross-sectional cohort, Gradient Tree Boosting machine learning models were trained and applied to predict the RSV infection status of a subject using the composition of their nasal microbiota, where status was defined in three ways: RSV infected vs. healthy; healthy vs. mild RSV infection vs. severe RSV infection; and severity score (with all healthy subjects having a score of 0). Five-fold cross-validation was employed, with 20% of samples being held out during training and then used to test the accuracy of the trained model. This approach can indicate how much information about a subject’s status is reflected in the composition of their nasal microbiota. Performance was best when distinguishing infected from healthy, which could be done with 90% accuracy using compositional profiles at the level of exact sequence variants. Distinguishing healthy, mild disease, and severe disease was less effective, with an accuracy of 77% being achieved using compositional profiles summarized based on taxonomic assignment at the level of species. Prediction of the continuous valued severity score exhibited the worst performance, with 38% of the variance in severity score being explained by the model, also using species level compositional profiles. Sequence variants classified as *Moraxella*, *Staphylococcus*, *Corynebacterium*, or *Streptococcus* comprised the top three most informative features across all three models. Model result summaries are in **Supplemental Tables 5-7**.

## Discussion

In this study, we characterize signatures of dysbiosis associated with RSV infection and illness severity in infant nasal microbiota. We identify differences in measures of microbial diversity and in the abundance of specific bacterial taxa between infants who develop RSV infections and those who don’t, and show that these differences manifest longitudinally before, during, and after illness in a number of distinct patterns. While these associations are consistent with observations made previously of nasopharyngeal microbiota during acute illness [13, 26], the findings reported here elucidate the temporal sequence and persistence of these phenomena beyond the period of acute illness, and demonstrate their occurrence in the nasal cavity.

Based on the observed patterns of differential abundance before, during, and after illness, the relationships between most of the taxa identified as significant and RSV infection may be assigned to one of three general categories. The first category includes taxa which change during and after illness relative to healthy controls of the same age. The dynamics of these taxa are consistent with RSV infection influencing the abundance of certain microbes; normal flora which dramatically diminish in a persistent way as a result of infection (Clostridia, Bacilli, & *Staphylococcus*). The second category consists of taxa increased in abundance in infants who develop RSV infection at all timepoints: before, during, and after illness. The patterns of occurrence of these taxa imply that they either are indicative of underlying factors making the host more susceptible to infection or directly contribute to infection susceptibility (Alphaproteobacteria, Gammaproteobacteria, Pseudomondales, and *Moraxella*). The third category is comprised of taxa which are significantly elevated in the healthy control subjects at the initial visit and the timepoint corresponding to illness. Such taxa (*Corynebacterium* and *Anaerococcus*) likely either reflect underlying protective qualities of the host, promote such qualities, or are directly protective themselves. The remaining three taxa identified as significant each exhibit unique patterns of differential abundance between groups and don’t fit well into any of the three categories. The interpretations of their associations with RSV infection are less clear, but the occurrence patterns of Betaproteobacteria and Haemophilus could be explained by an opportunistic or synergistic relationship, wherein RSV infection produces circumstances conducive to their increasing in abundance, while their increased abundance may contribute to infection severity. The occurrence patterns of *Gluconacetobacter* suggest that it may reflect or promote infection susceptibility, but convalescence corresponds to an unfavorable host environment and it diminishes in abundance.

Most RSV associated taxa are associated both with the presence or absence of infection, and also with disease severity. This implies that the biological underpinnings of these associations exist to varying degrees as opposed to being categorically distinct, and that this variation is reflected in illness severity during infection. However, the fact that our infected vs. healthy classifier outperforms our severe illness vs. mild illness vs. healthy classifier and the severity score regressor, and the fact that all significant taxa are associated with illness while only eight of them are associated with severity, suggest that the composition of nasal microbiota is more strongly associated with the difference between RSV infected and uninfected than it is with continuous variation along the gradient from health to severe illness.

The taxa which are differentially abundant at one month of age – prior to infection and illness – present intriguing possibilities. At a basic level, it may be possible to predict an infant’s risk of RSV infection in the first year of life based on the presence and abundance of these taxa at one month. Furthermore, understanding the mechanism by which these bacteria are associated with infection risk could provide valuable insights into immunological development or mucosal function. More speculatively, the possibility exists that the association is causal, which would suggest that these taxa may be suitable targets for prebiotic, probiotic, or antimicrobial interventions. Similar reasoning could be applied to *H. influenzae* and Betaproteobacteria Burkholderiales, which are not differentially abundant prior to illness but are associated with illness severity, and which could be targeted or assayed during infection to mitigate or predict severity. Whether these microbes merely reflect underlying factors that influence infection susceptibility, severity, and resistance, or contribute to them directly, the clinical significance of RSV infection in the short term, and respiratory infection-associated asthma and atopy in the long term, make these bacteria and their relationship to respiratory health important targets for translationally-oriented study.

We recognize a number of limitations of this study. Notably, our longitudinal cohort was substantially smaller than our cross-sectional cohort and incidentally it only contained two severely ill subjects. This prevented us from assessing associations with severity at the pre- and post-illness timepoints, which would be desirable. We were also limited to short read amplicon sequencing to profile the bacterial communities that were sampled. This inherently limits our ability to resolve species and strains of bacteria. Furthermore, marker gene assays contain no functional information about the microbial communities and no immunological information about the host. More comprehensive assays such as shotgun metagenomic sequencing and flow cytometry would greatly enrich our understanding of the systems of interest. Finally, all of our subjects were less than one year of age, had not been previously infected with RSV, and no subject was sampled more than approximately one month after illness, which prevented us from examining microbiota-RSV associations among infants who became infected in their second year of life or who had recurrent infections, and made it impossible to determine how long the associations we observed persisted after illness. Similarly, our earliest samples were at approximately one month of age and already showed differences between subjects who went on to acquire RSV infections and those who did not, so we were unable to determine how early those differences manifested. Nevertheless, our findings provide novel insight into the developmental dynamics of the nasal microbiome in the first year of life as they relate to susceptibility, acute illness, severity, and convalescence associated with first-time RSV infection.

## Availability of data and materials

All phenotypic data, 16S rRNA sequence reads and generated datasets is publicly available through dbGaP accession phs001201.v2.p1.

## Author Contributions

S.R.G., E.E.W., and M.T.C. conceptualized the study. S.R.G., E.E.W., T.J.M., M.T.C., and A.G. designed the experiments. E.E.W., M.T.C., and A.R.F. developed the cohort, and collected the specimens and clinical data. J.H.-W., S.B., J.J., and A.C. facilitated data organization, management and analysis. T.J.M., M.T.C., E.E.W., A.L.G., H.A.K., S.R.G., J.H.W., J.J., S.B., A.C., and A.G. generated, analyzed and interpreted the data. S.R.G. and A.G. wrote and/or revised the manuscript. All authors read and approved the final manuscript.

## Acknowledgements

We thank Amy Murphy R.N., Mary Criddle, R.N., and Doreen Francis R.N. for assistance in recruiting and following study subjects. Microbiome sequencing in this study was completed by the University of Rochester Genomics Research Center (GRC).

## Supplemental Methods

### Genomic DNA extraction

Total genomic DNA was extracted from the nasal samples using a modification of the ZymoBIOMICS™ DNA Miniprep Kit (Zymo Research, Irvine, CA) and FastPrep mechanical lysis (MPBio, Solon, OH). 16S ribosomal DNA (rRNA) was amplified with Phusion High-Fidelity polymerase (Thermo Scientific, Waltham, MA) and dual indexed primers specific to the V1-V3 (8F: 5’ AGAGTTTGATCCTGGCTCAG 3’; 534R: 3’ ATTACCGCGGCTGCTGG 5’) hypervariable regions [1]. Amplicons were pooled and paired-end sequenced on an Illumina MiSeq (Illumina, San Diego, CA) in the University of Rochester Genomics Research Center. Each sequencing run included: (1) positive controls consisting of a 1:5 mixture of *Staphylococcus aureus*, *Lactococcus lactis*, *Porphyromonas gingivalis*, *Streptococcus mutans*, and *Escherichia coli*; and (2) negative controls consisting of sterile saline.

### Microbiota background control

The background microbiota was monitored at multiple stages of sample collection and processing. All sterile saline, buffers, reagents, plasticware and flocked nylon swabs used for sample collection, extraction and amplification of nucleic acid were UV irradiated to eliminate possible DNA background contamination. Elimination of potential background from the irradiated buffers, reagents, plasticware and swabs was confirmed by 16S rRNA amplification. After sample collection, multiple aliquots of sterile saline with swabs used for sample collection were carried through our entire sequencing protocol as individual samples, including DNA extraction, 16S rRNA amplification, library construction and sequencing to monitor potential background microbiome [2]. Data from these background control samples is deposited in SRA along with positive controls.

### Bioinformatics analysis

Raw data from the Illumina MiSeq was first converted into FASTQ format 2×312 paired end sequence files using the bcl2fastq program, version 1.8.4, provided by Illumina. Format conversion was performed without de-multiplexing and the EAMMS algorithm was disabled. All other settings were default. Reads were multiplexed using a configuration described previously [1]. Briefly, for both reads in a pair, the first 12 bases were a barcode, which was followed by a primer, then a heterogeneity spacer, and then the target 16S rRNA sequence. QIIME 1.9.1 [3] was used to extract the barcodes into a separate file for importing into QIIME 2 [4], which was used to perform all subsequent processing. Reads were demultiplexed requiring exact barcode matches, and 16S primers were removed allowing 20% mismatches and requiring at least 18 bases. Cleaning, joining, and denoising were performed using DADA2 [5]: forward reads were truncated to 275 bps and reverse reads to 260 bps, error profiles were learned with a sample of one million reads, and a maximum expected error of two was allowed. Taxonomic classification was performed with a custom naïve Bayesian classifier trained on the August, 2013 release of GreenGenes [6, 7]. Sequence variants that could not be classified at least at the phylum level were discarded. Sequencing variants observed fewer than ten times total, or in only one sample, were discarded. Samples with fewer than 900 reads were discarded.

Phylogenetic trees were constructed for each cohort using MAFFT for sequence alignment and FastTree for tree construction [8, 9]. Prior to diversity analyses, samples were rarefied to a depth of 900 reads. Faith’s PD and the Shannon index were used to measure alpha diversity, and Kruskal-Wallis to test for differences. Weighted and Unweighted Unifrac distances were used to measure beta diversity [10] and pairwise PERMANOVA to test for differences.

Infected vs. healthy and healthy vs. mild vs. severe classification, and severity score regression, were performed using the Sample Classifier plugin [11] in QIIME 2, using the Gradient Tree Boosting Classifier/Regressor, five-fold cross-validation, 20% data hold-out for testing, 5,000 estimators, parameter tuning, and feature selection. Both exact sequence variant abundances and abundances of taxa summarized at species level were tried as inputs, and whichever performed better was used and reported.

### Associations of taxon abundance with RSV infection and disease severity

Univariate tests for differential taxon abundance between groups was performed using both ANCOM [12] and LefSe [13]. Multivariate regression models using gneiss [14] and MaAsLin [15] were employed to assess associations of taxon abundance with RSV infection and disease severity while controlling for the potentially confounding covariates sex, mode of delivery, age at sampling, reads per sample, and antibiotic usage. The cross-sectional and longitudinal cohorts were analyzed independently. All reported results were significant by at least two tests.

## References

1. Clemente JC, Ursell LK, Parfrey LW, Knight R. The impact of the gut microbiota on human health: an integrative view. Cell 2012; 148:1258–70.

2. Caporaso JG, Lauber CL, Costello EK, et al. Moving pictures of the human microbiome. Genome biology 2011; 12:R50.

3. Faust K, Sathirapongsasuti JF, Izard J, et al. Microbial co-occurrence relationships in the human microbiome. PLoS Comput Biol 2012; 8:e1002606.

4. Grier A, McDavid A, Wang B, et al. Neonatal gut and respiratory microbiota: coordinated development through time and space. Microbiome 2018; 6:193.

5. Hofstra JJ, Matamoros S, van de Pol MA, et al. Changes in microbiota during experimental human Rhinovirus infection. BMC Infect Dis 2015; 15:336.

6. Xu Q, Wischmeyer J, Gonzalez E, Pichichero ME. Nasopharyngeal polymicrobial colonization during health, viral upper respiratory infection and upper respiratory bacterial infection. The Journal of infection 2017; 75:26–34.

7. Blander JM, Longman RS, Iliev ID, Sonnenberg GF, Artis D. Regulation of inflammation by microbiota interactions with the host. Nat Immunol 2017; 18:851–60.

8. Gollwitzer ES, Saglani S, Trompette A, et al. Lung microbiota promotes tolerance to allergens in neonates via PD-L1. Nature medicine 2014; 20:642–7.

9. Olszak T, An D, Zeissig S, et al. Microbial exposure during early life has persistent effects on natural killer T cell function. Science 2012; 336:489–93.

10. Hooper LV, Littman DR, Macpherson AJ. Interactions between the microbiota and the immune system. Science 2012; 336:1268–73.

11. Man WH, de Steenhuijsen Piters WA, Bogaert D. The microbiota of the respiratory tract: gatekeeper to respiratory health. Nature reviews Microbiology 2017; 15:259–70.

12. Bosch A, de Steenhuijsen Piters WAA, van Houten MA, et al. Maturation of the Infant Respiratory Microbiota, Environmental Drivers, and Health Consequences. A Prospective Cohort Study. Am J Respir Crit Care Med 2017; 196:1582–90.

13. de Steenhuijsen Piters WA, Heinonen S, Hasrat R, et al. Nasopharyngeal Microbiota, Host Transcriptome, and Disease Severity in Children with Respiratory Syncytial Virus Infection. Am J Respir Crit Care Med 2016; 194:1104–15.

14. de Steenhuijsen Piters WA, Sanders EA, Bogaert D. The role of the local microbial ecosystem in respiratory health and disease. Philos Trans R Soc Lond B Biol Sci 2015; 370.

15. Hilty M, Qi W, Brugger SD, et al. Nasopharyngeal microbiota in infants with acute otitis media. J Infect Dis 2012; 205:1048–55.

16. Pettigrew MM, Laufer AS, Gent JF, Kong Y, Fennie KP, Metlay JP. Upper respiratory tract microbial communities, acute otitis media pathogens, and antibiotic use in healthy and sick children. Applied and environmental microbiology 2012; 78:6262–70.

17. Hall CB, Weinberg GA, Blumkin AK, et al. Respiratory syncytial virus-associated hospitalizations among children less than 24 months of age. Pediatrics 2013; 132:e341–8.

18. Stockman LJ, Curns AT, Anderson LJ, Fischer-Langley G. Respiratory syncytial virus-associated hospitalizations among infants and young children in the United States, 1997-2006. Pediatr Infect Dis J 2012; 31:5–9.

19. Tregoning JS, Schwarze J. Respiratory viral infections in infants: causes, clinical symptoms, virology, and immunology. Clin Microbiol Rev 2010; 23:74–98.

20. Boyce TG, Mellen BG, Mitchel EF, Jr., Wright PF, Griffin MR. Rates of hospitalization for respiratory syncytial virus infection among children in medicaid. The Journal of pediatrics 2000; 137:865–70.

21. Garcia CG, Bhore R, Soriano-Fallas A, et al. Risk factors in children hospitalized with RSV bronchiolitis versus non-RSV bronchiolitis. Pediatrics 2010; 126:e1453–60.

22. Glezen WP, Taber LH, Frank AL, Kasel JA. Risk of primary infection and reinfection with respiratory syncytial virus. Am J Dis Child 1986; 140:543–6.

23. Proud D, Chow CW. Role of viral infections in asthma and chronic obstructive pulmonary disease. Am J Respir Cell Mol Biol 2006; 35:513–8.

24. Fonseca W, Lukacs NW, Ptaschinski C. Factors Affecting the Immunity to Respiratory Syncytial Virus: From Epigenetics to Microbiome. Front Immunol 2018; 9:226.

25. Teo SM, Mok D, Pham K, et al. The infant nasopharyngeal microbiome impacts severity of lower respiratory infection and risk of asthma development. Cell host & microbe 2015; 17:704–15.

26. Rosas-Salazar C, Shilts MH, Tovchigrechko A, et al. Differences in the Nasopharyngeal Microbiome During Acute Respiratory Tract Infection With Human Rhinovirus and Respiratory Syncytial Virus in Infancy. J Infect Dis 2016; 214:1924–8.

27. Biesbroek G, Tsivtsivadze E, Sanders EA, et al. Early respiratory microbiota composition determines bacterial succession patterns and respiratory health in children. Am J Respir Crit Care Med 2014; 190:1283–92.

28. Lynch JP, Sikder MA, Curren BF, et al. The Influence of the Microbiome on Early-Life Severe Viral Lower Respiratory Infections and Asthma-Food for Thought? Front Immunol 2017; 8:156.

29. Walsh EE, Mariani TJ, Chu C, et al. Aims, Study Design, and Enrollment Results From the Assessing Predictors of Infant Respiratory Syncytial Virus Effects and Severity Study. JMIR Res Protoc 2019; 8:e12907.

30. Bolyen E, Rideout JR, Dillon MR, et al. Reproducible, interactive, scalable and extensible microbiome data science using QIIME 2. Nat Biotechnol 2019; 37:852–7.

31. Callahan BJ, McMurdie PJ, Rosen MJ, Han AW, Johnson AJ, Holmes SP. DADA2: High-resolution sample inference from Illumina amplicon data. Nature methods 2016; 13:581–3.

32. DeSantis TZ, Hugenholtz P, Larsen N, et al. Greengenes, a chimera-checked 16S rRNA gene database and workbench compatible with ARB. Applied and environmental microbiology 2006; 72:5069–72.

33. McDonald D, Price MN, Goodrich J, et al. An improved Greengenes taxonomy with explicit ranks for ecological and evolutionary analyses of bacteria and archaea. The ISME journal 2012; 6:610–8.

34. Hawinkel S, Mattiello F, Bijnens L, Thas O. A broken promise: microbiome differential abundance methods do not control the false discovery rate. Brief Bioinform 2019; 20:210–21.

35. Weiss S, Xu ZZ, Peddada S, et al. Normalization and microbial differential abundance strategies depend upon data characteristics. Microbiome 2017; 5:27.

36. Caserta MT, Qiu X, Tesini B, et al. Development of a Global Respiratory Severity Score for Respiratory Syncytial Virus Infection in Infants. J Infect Dis 2017; 215:750–6.

## References

1. Fadrosh DW, Ma B, Gajer P, et al. An improved dual-indexing approach for multiplexed 16S rRNA gene sequencing on the Illumina MiSeq platform. Microbiome 2014; 2:6.

2. Grier A, McDavid A, Wang B, et al. Neonatal gut and respiratory microbiota: coordinated development through time and space. Microbiome 2018; 6:193.

3. Caporaso JG, Kuczynski J, Stombaugh J, et al. QIIME allows analysis of high-throughput community sequencing data. Nature methods 2010; 7:335–6.

4. Bolyen E, Rideout JR, Dillon MR, et al. Reproducible, interactive, scalable and extensible microbiome data science using QIIME 2. Nat Biotechnol 2019; 37:852–7.

5. Callahan BJ, McMurdie PJ, Rosen MJ, Han AW, Johnson AJ, Holmes SP. DADA2: High-resolution sample inference from Illumina amplicon data. Nature methods 2016; 13:581–3.

6. DeSantis TZ, Hugenholtz P, Larsen N, et al. Greengenes, a chimera-checked 16S rRNA gene database and workbench compatible with ARB. Applied and environmental microbiology 2006; 72:5069–72.

7. McDonald D, Price MN, Goodrich J, et al. An improved Greengenes taxonomy with explicit ranks for ecological and evolutionary analyses of bacteria and archaea. The ISME journal 2012; 6:610–8.

8. Katoh K, Standley DM. MAFFT multiple sequence alignment software version 7: improvements in performance and usability. Mol Biol Evol 2013; 30:772–80.

9. Price MN, Dehal PS, Arkin AP. FastTree 2--approximately maximum-likelihood trees for large alignments. PloS one 2010; 5:e9490.

10. Lozupone C, Lladser ME, Knights D, Stombaugh J, Knight R. UniFrac: an effective distance metric for microbial community comparison. The ISME journal 2011; 5:169–72.

11. Bokulich NA, Dillon MR, Zhang Y, et al. q2-longitudinal: Longitudinal and Paired-Sample Analyses of Microbiome Data. mSystems 2018; 3.

12. Mandal S, Van Treuren W, White RA, Eggesbo M, Knight R, Peddada SD. Analysis of composition of microbiomes: a novel method for studying microbial composition. Microb Ecol Health Dis 2015; 26:27663.

13. Segata N, Izard J, Waldron L, et al. Metagenomic biomarker discovery and explanation. Genome biology 2011; 12:R60.

14. Morton JT, Sanders J, Quinn RA, et al. Balance Trees Reveal Microbial Niche Differentiation. mSystems 2017; 2.

15. Morgan XC, Tickle TL, Sokol H, et al. Dysfunction of the intestinal microbiome in inflammatory bowel disease and treatment. Genome biology 2012; 13:R79.

